# Target of Rapamycin Complex 2 modulates development through Hedgehog/Patched signaling in *C. elegans*

**DOI:** 10.1101/2022.11.17.517002

**Authors:** Sinclair W. Emans, Armen Yerevanian, Fasih M. Ahsan, Yifei Zhou, Lucydalila Cedillo, Alexander A. Soukas

## Abstract

Both Hedgehog (Hh) signaling and target of rapamycin complex 2 (TORC2) are central, evolutionarily conserved pathways that regulate development and metabolism. In *C. elegans*, loss of essential TORC2 component RICTOR (*rict-1*) causes delayed development, shortened lifespan, reduced brood, small size, and increased fat. Here we report that knockdown of Hedgehog-related morphogen *grd-1* and its Patched-related receptor *ptr-11* rescues delayed development in TORC2 loss of function mutants, indicating an unexpected role for *grd-1*/*ptr-11* in slowing developmental rate downstream of nutrient sensing pathways. Further, we implicate chronic stress transcription factor *pqm-1* as a key transcriptional effector of *grd-1*/*ptr-11* in slowing whole-organism growth. We propose that the TORC2/*grd-1*/*ptr-11*/*pqm-1* signaling relay acts as a critical executor of growth to slow development when *C. elegans* encounters unfavorable growth conditions.

**Summary statement:** Developmental rate in *C. elegans* is dramatically slowed in animals deficient in nutrient-sensitive target of rapamycin complex 2 signaling and slowing is effected by increased activity of a previously uncharacterized Hh-r/Ptr signaling relay.

## INTRODUCTION

Hedgehog (Hh) morphogens are highly conserved positive regulators of development found throughout vertebrates and invertebrates (Aspöck et al., 1999; Echelard et al., 1993; Nüsslein-Volhard and Wieschaus, 1980). Extensive study in organisms such as *Drosophila melanogaster* has revealed a canonical pathway wherein Hh proteins bind to Patched (Ptc) allowing for shifts in gene expression via the Gli transcription factors (Ingham, 2022). However, these proteins remain relatively unexplored in the model organism *Caenorhabditis elegans*. In *C. elegans*, divergence from the canonical signaling pathway leaves some Hh proteins missing while expanding the Hh-related (Hh-r) and Patched-related (Ptr) protein families from just 5 members to over 80 members (Aspöck et al., 1999). Of these, only 14 have been mechanistically characterized and just two, *wrt-10* and *grl-21*, have been functionally associated with Patched/Patched-related receptors (Lin and Wang, 2017; Templeman et al., 2020). From molting to reproductive aging, all of the Hh-r proteins that have been studied serve important, non-redundant roles in *C. elegans*. Hh-r proteins such as *qua-1* are indispensable to larval transition while others such as *wrt-10* govern aspects of healthspan downstream of important, conserved regulators such as CREB (Hao et al., 2006; Templeman et al., 2020).

Activity of highly conserved, nutrient-sensing, signaling pathways is integral to ensuring normal growth rate in *C. elegans*. For instance, reduced target of rapamycin complex 2 (TORC2) signaling has pronounced effects on both developmental rate (Jones et al., 2009; Soukas et al., 2009). Loss of function of the gene encoding the essential TORC2 subunit *rictor* extends time to adulthood at 20ºC from 48 hrs. to 72 hrs., reduces size, lowers brood, increases fat, and shortens lifespan (Jones et al., 2009; Soukas et al., 2009). Our previous work showed that suppression of essential dosage compensation complex (DCC) member *dpy-21* downstream of the TORC2 effector kinase serum- and glucocorticoid-induced kinase 1 (SGK-1) was sufficient to rescue the delayed development, reduced brood, and increased fat of *rict-1* mutants – but not their reduced body size and shortened lifespan – via action of histone methyltransferases SET-1 and SET-4 (Webster et al., 2013). However, the spectrum of effectors of TORC2 governance over developmental rate and healthy growth remain incompletely characterized.

In this study, we identify a *grd-1*/*ptr-11* signaling relay that negatively regulates development downstream of TORC2. We find that, like many other Hh-r proteins, *grd-1* expression is animated during molting transitions, and, surprisingly, *grd-1* knockdown by RNAi leads to developmental acceleration in wild type animals. Further, *grd-1* knockdown significantly rescues TORC2 mutants’ slowed development and TORC2 signaling likely modulates *grd-1* activity rather than expression. We also identify *ptr-11* as a likely recipient of *grd-1* signaling. *ptr-11* knockdown phenocopies the ability of *grd-1* knockdown to accelerate growth in wild type animals and to restore normal growth rate in TORC2 mutants. Importantly, augmented *grd-1* expression is sufficient to delay development, and *ptr-11* knockdown significantly rescues the delayed development of *grd-1* overexpressor animals. Finally, we identify *pqm-1* as a potential effector of the *grd-1*/*ptr-11* signaling cascade. *pqm-1* knockdown partially phenocopies *grd-1*/*ptr-11* knockdown and *pqm-1* target genes are upregulated in TORC2 loss of function mutants in a manner dependent upon both *grd-1* and *ptr-11*. In aggregate, we propose a TORC2/SGK-1/GRD-1/PTR-11/PQM-1 signaling relay as the effector arm by which TORC2 slows development in response to unfavorable environmental conditions in *C. elegans*.

## MATERIALS AND METHODS

### Strains and maintenance

C. elegans animals were grown and maintained at 20°C on Nematode Growth Media (NGM) seeded with *Escherichia coli* OP50-1 as previously described (Soukas et al., 2009). The following strains were used: wild type N2 (Bristol), MGH266 *rict-1*(*mg451*), MGH300 *sgk-1*(*mg455*), CB1370 *daf-2*(*e1370*), DA465 *eat-2*(*ad465*), VC222 *raga-1*(*ok386*), MGH9 *rsks-1*(*ok1255*), MGH27 *rict-1*(*mg451*);*sgk-1*(*mg455*), MGH35 *sinh-1*(*mg452*), MGH629 *ptr-11*::*GFP*::*AID\**::*3xFLAG*::*ptr-11 3’UTR*, MGH531 *grd-1p*::*grd-1g*::*grd-1 3’UTR*, MGH563 *grd-1p*::*GFP*, SYS573 *pqm-1p*::*GFP*::*pqm-1*, and MGH618 *rict-1*(*mg451*);*pqm-1p*::*GFP*::*pqm-1*.

Animals were synchronized by hypochlorite bleach treatment of a population of gravid adults. Animals were collected in M9 medium, centrifuged at 4400 rpm for 1 minute and resuspended in 6 mL 1.3% bleach, 250 mM NaOH for 1 minute of vigorous shaking. Bleach solution exposure and shaking step was repeated after one wash in M9. After the second bleach step, pellet was resuspended in minimal M9 by gentle pipetting, and washed in M9 four times. Eggs were resuspended in 12 mL M9, and solution was left to rotate overnight for 23 hours at 20°C.

### RNA interference (RNAi)

RNAi plates were prepared with standard NGM media mixed with 5 mM isopropyl-B-D-thiogalactopyranoside and 200 ug/mL carbenicillin. All RNAi clones were isolated from the genome wide *E. coli* HT115 Ahringer library (Horizon Discovery) and sequence verified before use. RNAi clones were obtained by seeding onto ampicillin/tetracycline treated Luria Broth (LB) plates. RNAi clones were grown overnight at 37°C with shaking for 18 hours in LB with 200 ug/mL carbenicillin. Cultures were spun down at 4400 rpm for 15 minutes, pellets resuspended in one tenth starting LB volume and dispensed onto RNAi plates no more than 48 hours prior to adding worms.

### Developmental timing assays

Synchronized L1 animals prepared as described above were dropped onto RNAi plates with vector RNAi control or RNAi directed against the gene of interest at 20°C. Time to adulthood was measuring by evaluating the appearance of the vulvar slit in hermaphrodite animals. Two to three biological replicates were performed for all conditions and data is available in supplementary Table S1.

Developmental screening was performed by evaluating relative proportion of wild type adults on treatment RNAi as compared to vector RNAi at 49 hours. Three biological replicates were pooled for analysis.

### Brood size assay

Synchronized L1 animals were dropped onto RNAi plates with the appropriate RNAi clone. After having reached young adult (YA) stage, two hermaphrodites were transferred onto five plates with the corresponding RNAi clone for a total of ten adults per condition. Adults were transferred every day for five days. Brood was measured as the total amount of progeny on the plates after three days at 20°C.

### Body fat mass measurement

At day one of adulthood, synchronous animals were collected in M9, washed once, centrifuged at 500 rpm for 1 minute, and resuspended in 40% isopropanol for 3 minutes. Fixed worms were then stained with 3 ug/mL Nile red in 40% isopropanol for 2 hours. Animals were collected in M9 supplemented with 0.01% Triton-X100, mounted on glass slides, and imaged in the GFP channel for 10 ms at 5x magnification (Pino et al., 2013). Body fat mass was measured as the fluorescence intensity relative to body area in pixels using ImageJ.

### Body size measurement

Adult day one animals were mounted on 2% agarose pads in 2.5 mM levamisole and imaged by brightfield microscopy at 2.5x. Body area was determined in pixels using ImageJ.

### Longevity assay

Synchronous L1 animals were dropped onto the appropriate RNAi clones and allowed to grow to L4/YA stage. 40-60 worms were then transferred to four plates with the corresponding RNAi clone supplemented with 10-50 uM 5-fluorodeoxyuridine (FUDR) for a total of ∼200 adults per condition. Animals were scored as dead or alive by movement or response to gentle prodding every other day. Data were analyzed using OASIS2 software package (https://sbi.postech.ac.kr/oasis2/).

### Quantitative RT-PCR

Worms were collected in TRIzol Reagent (Invitrogen), flash frozen in liquid nitrogen, and stored at -80°C prior to RNA isolation. Samples were lysed using metal beads and the Tissuelyser I system (Qiagen).

RNA was isolated from lysate using Direct-zol RNA Miniprep kit (Zymo). cDNA samples were synthesized with the QuantiTect reverse transcription kit (Qiagen). Quantitative RT-PCR was performed with the QuantiTect SYBR Green RT-PCR kit (Qiagen) in a Bio-Rad CFX96 RT-PCR thermocycler. Fold changes were determined via the 2^-^Δ Δ _Ct_ method as previously described (Livak and Schmittgen, 2001). Primers used:

*mlt-10* F: GGCCTTGGCAGCCGTAAC

*mlt-10* R: TAAGCTCCACGGATGAGGTC

*grd-1* F: CCGCTCTGCTGATATAAACCACG

*grd-1* R: TCATCGCAACATTCTACCGT

*ptr-11* F: AGCCGCCTATCCGGTTTATT

*ptr-11* R: GACACGGGTTTCATATCCAGC

*C49G7*.*7* F: CGGAGATCGGGAAACCCTTT

*C49G7*.*7* R: GGGCGGCAAGGAAAGTTAAA

*F18E3*.*12* F: CAATTTCCACCACACCAGCC

*F18E3*.*12* R: TGAGCGCAGTTGAAATCGTTG

*ugt-43* F: AGTTACCGGTCATTCTCATTTAAAGTT

*ugt-43* R: TGAGTGGTAAGAGAAGAGTCACA

*C15B12*.*8* F: TTGTGACGATCCCAGGAAGC

*C15B12*.*8* R: GTCCGGGGAGTTGGTGTATT

*F33H12*.*7* F: TGGATTTTTGGAACACAAACGA

*F33H12*.*7* R: CGCACCGGAAAGGTCTACTT

*str-7* F: ACGCGTTTTTCGGTTTTATCCT

*str-7* R: GAGGTGGAGGAACGTGTGAA

### Microscopy

All imaging unless otherwise specified was performed by mounting worms on 2% agarose pads in 2.5 mM levamisole using the Leica THUNDER Imager system. Imaging was performed within 5 minutes of sample preparation. Binning measurements were done with a minimum population of 10 worms per replicate. *grd-1p*::*GFP* activation was defined as intestinal expression.

### Molting midpoint analysis

The molting midpoints were defined as the times at which ∼50% of the worm population is molting and were determined as the midpoint between two minima in *mlt-10* expression as shown previously (McCulloch and Rougvie, 2014).

### Statistical analysis

All statistical analyses and representations were performed in either Prism 9 (Graphpad) or Bioconductor (R). LogEC50 for developmental curves was determined by fitting a variable slope sigmoidal dose-response curve to each curve. The difference between the vector and *grd-1* RNAi curves was calculated by percent difference between the logEC50 of each curve.

### RNA sequencing analysis

L3 animals treated with empty vector (EV) control or *grd-1* RNAi (*grd-1*) were collected in paired, biologically independent triplicate experiments. RNA was collected and purified in the same way as described above for qRT-PCR. Total purified RNA samples were sent to Azenta (Genewiz) for quality control, library preparation, and mRNA sequencing. Samples were first verified for RNA integrity scores greater than 8 using an Agilent Tapestation 4200. Illumina library preparation was performed using polyA selection for mRNA species. Approximately 20 million paired-end, 150 base pair reads were obtained per sample.

Read filtering and quasi-alignment were performed using custom UNIX/bash shell scripts on the Mass General Brigham ERISOne Scientific Computing Linux Cluster. Reads were analyzed for quality control using FastQC (https://www.bioinformatics.babraham.ac.uk/projects/fastqc/) and MultiQC (Ewels et al., 2016), and filtered for adapter contamination, truncated short reads, or low-quality bases using BBDuk (Grigoriev et al., 2012). Trimmed, cleaned reads were then quantified against the *C. elegans* reference transcriptome annotation (WBcel235, Ensembl Release 105) using Salmon, correcting for sequencing and GC content bias using the command parameters “--seqBias” and “—gcBias”, respectively (Cunningham et al., 2022; Patro et al., 2017).

All statistical analysis and visualizations were performed using the R (https://www.r-project.org) Bioconductor (http://www.nature.com/nmeth/journal/v12/n2/abs/nmeth.3252.html) environment. Quasi-aligned transcript quantification files for each sample were collapsed into gene-level count matrices using R package tximport (Soneson et al., 2015), and paired differential expression was calculated using R package DESeq2 (Love et al., 2014) with a design formula of “∼Replicate + Treatment”, where ‘Replicate’ accounts for inter-replicate batch effect variation in the paired experimental samples. Genes were considered differentially expressed with a Benjamini-Hochberg False Discovery Rate (FDR) corrected *P* value < 0.05 and an absolute log2 transformed fold change of 1.5 (Benjamini and Hochberg, 1995).

The top 100 genes contributing to variation in principal component 1 (PC1 – explanatory for the divergence seen between EV and grd-1 RNAi) were extracted and assessed for Gene Ontology (GO) term pathway overrepresentation using R package clusterProfiler (Wu et al., 2021). To perform transcription factor target enrichment analyses, promoter sequences 1500 bases upstream and 500 bases downstream of the transcription start site of significantly upregulated and downregulated genes were extracted using R package TxDb.Celegans.UCSC.ens11.ensGene (https://doi.org/doi:10.18129/B9.bioc.TxDb.Celegans.UCSC.ce11.ensGene). Transcription factor binding sites were retrieved from the modENCODE modMine v33 database (Contrino et al., 2012) and filtered for presence in the promoter sequences of the differentially expressed gene sets. Total binding site ratios in both upregulated and downregulated genes were tallied (defined as the number of binding sites identified for a given transcription factor divided by all transcription factor binding site identified) and visualized using Prism 9 (Graphpad). Significance of the enrichment in observed binding site ratio versus expected ratio was calculated using a two-tailed exact binomial test with a 95% confidence level using R function binom_test(), and FDR adjusted *P* values were obtained using R function p.adjust().

## RESULTS

### Knockdown of Hedgehog-related morphogen *grd-1* unexpectedly accelerates *C. elegans* development

Knockdown of Hh-r proteins in *C. elegans* has been reported to prompt developmental delay, arrest, or defects (Cohen et al., 2021; Hao et al., 2006; Zugasti et al., 2005). Besides high-level transcriptional analyses, the Groundhog (GRD) family is absent from previous investigations into Hh-r proteins, and we sought to determine whether knockdown of these proteins affects worm development (Aspöck et al., 1999). We performed a small-scale screen of *grd* family RNAi on larval development, finding that although *grd-7* knockdown does slow larval development, *grd-1* knockdown accelerates development (Fig. 1A-B; Table S1). We confirmed that the Ahringer RNAi clone reduces *grd-1* transcript levels by ∼90% (Fig. S1A).

**Fig. 1.**
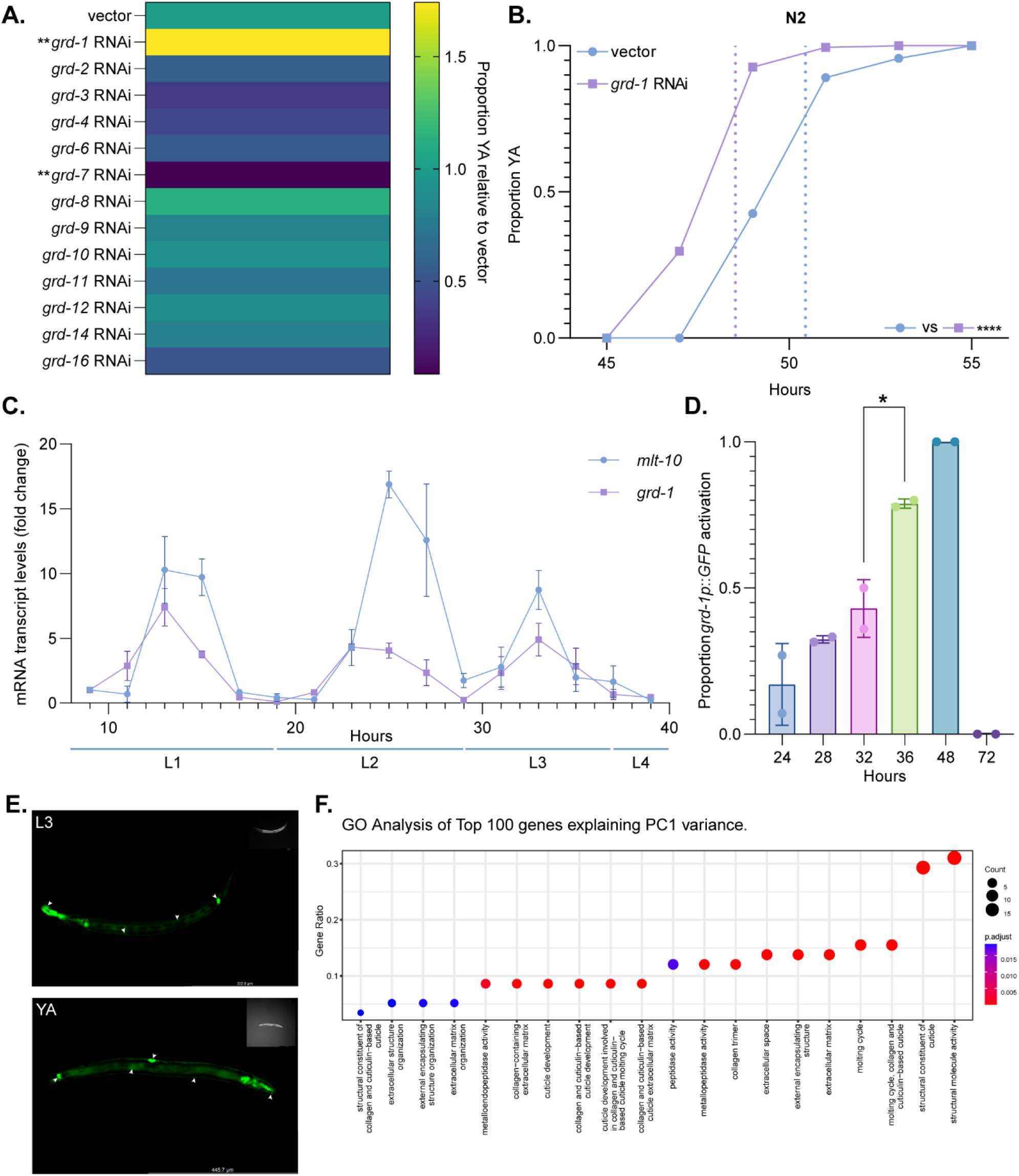
Knockdown of cyclically-expressed *grd-1* by RNAi accelerates wild type development. (**A**) Developmental screening of *grd* family members reveals that *grd-1* RNAi uniquely and unexpectedly accelerates wild type *C. elegans* development (*n* = 3, \*\**P*<0.05 by one-way ANOVA with two-stage Benjamini-Krieger-Yekutieli FDR-adjustment). Note that vector control = 1. (**B**) Developmental rate in wild type animals is accelerated by 2 hours on *grd-1* RNAi (*n* ≥ 100, \*\*\*\**P*<0.0001 by Bonferroni corrected log-rank test). Dashed lines represent hypothetical midpoint times at which 50% of the population has reached young adulthood. Data in biological triplicate in supplementary Table S1. (**C**) Measurement of mRNA transcript levels for *grd-1* and *mlt-10* from 9 hours to 39 hours reveals that *grd-1* is expressed cyclically with molting cycles. Fold changes normalized to vector at 9 hours. Data from three replicates. (**D**) *grd-1p*::*GFP* animals imaged from mid L3 to adult day 1 (AD1) show intestinal GFP expression that peaks at molting midpoints of 36 and 48 hours and is totally extinguished in day one of adulthood at 72 hours *(n* > 20, P<0.05 by one-way ANOVA with Dunnett’s correction for multiple comparisons). Data represented as mean ± s.e.m. (**E**) Representative images of an L3 larval animal with intestinal, hypodermal, head neuron, and rear epithelial *grd-1* expression (*top panel*) and a young adult (YA) exhibiting *grd-1* expression in the intestine, rear epithelial cells, vulva, hypodermis, and head neurons (*bottom panel*). (**F**) GO-term enrichment analysis of the top 100 genes contributing to principal component 1 (PC1) variance in RNA-sequencing of *grd-1* suppressed L3 animals shows the strongest signal for cuticle genes and other molting associated factors (*n* = 3, adjusted *P* < 0.05, see supplementary Table S2 for differentially expressed genes and top 100 most variance PC1 genes).

Previous transcriptomic profiling of *C. elegans* development indicates cyclical expression of *grd-1* mRNA (Hendriks et al., 2014), which we confirmed via qRT-PCR (Fig. 1C). Further, a *grd-1*::GFP promoter reporter confirms a significant increase in *grd-1* expression during the L3/L4 molt and the L4/young adult (YA) molt, specifically in the intestine (Fig. 1D). We also replicated previous reports of *grd-1* expression in rectal epithelial cells (Aspöck et al., 1999) and further observe expression in other tissues including head neurons, hypodermis, intestine, and vulva (Fig. 1E).

Having determined that *grd-1* is necessary for normal developmental rate in wild type animals and that it is cyclically expressed with molting, we performed an RNA sequencing analysis of *grd-1* RNAi vs. vector at the L3 developmental stage in order to ascertain whether molting factors are relevant to the acceleration caused by *grd-1* knockdown. Indeed, we find that analysis of the top 100 differentially expressed genes explaining the variance caused by *grd-1* knockdown indicates enrichment for GO terms matching molting genes (Fig. 1F; Table S2).

### *C. elegans* TORC2 mutant *rict-1* is sensitized to *grd-1* knockdown

Larval progression is gated by nutritional rheostats and heterochronic genes that tightly determine when and if the animal transitions from one larval stage to the next (Mata-Cabana et al., 2021; Moss, 2007). Based upon its developmental pattern of expression, we hypothesized that *grd-1* may act downstream of major controllers of larval growth. We assessed the effect of *grd-1* RNAi on the development of several mutants in growth factor/nutrient sensing pathways that manifest growth delay including *daf-2*(*e1370*) (insulin-like receptor hypomorph), *eat-2*(*ad465*) (defective pharyngeal pumping which leads to caloric restriction), *raga-1*(*ok386*) (hypomorphic defects in mTOR complex 1 signaling), *rsks-1*(*ok1255*) (S6 Kinase), and *rict-1*(*mg451*) (mTOR complex 2 loss of function) (Fig. 2A-E; Table S1). As in wild type, *grd-1* RNAi accelerates all the mutants’ L1 to young adult (YA) development time by ∼2 hours, except for *rict-1* mutants which matured ∼6 hours faster (Fig. 2E). Moreover, the proportional acceleration in median time to adulthood is significantly greater in *rict-1* mutants subjected to *grd-1* RNAi relative to acceleration seen in wild type and other mutants assessed (Fig. 2F).

**Fig. 2.**
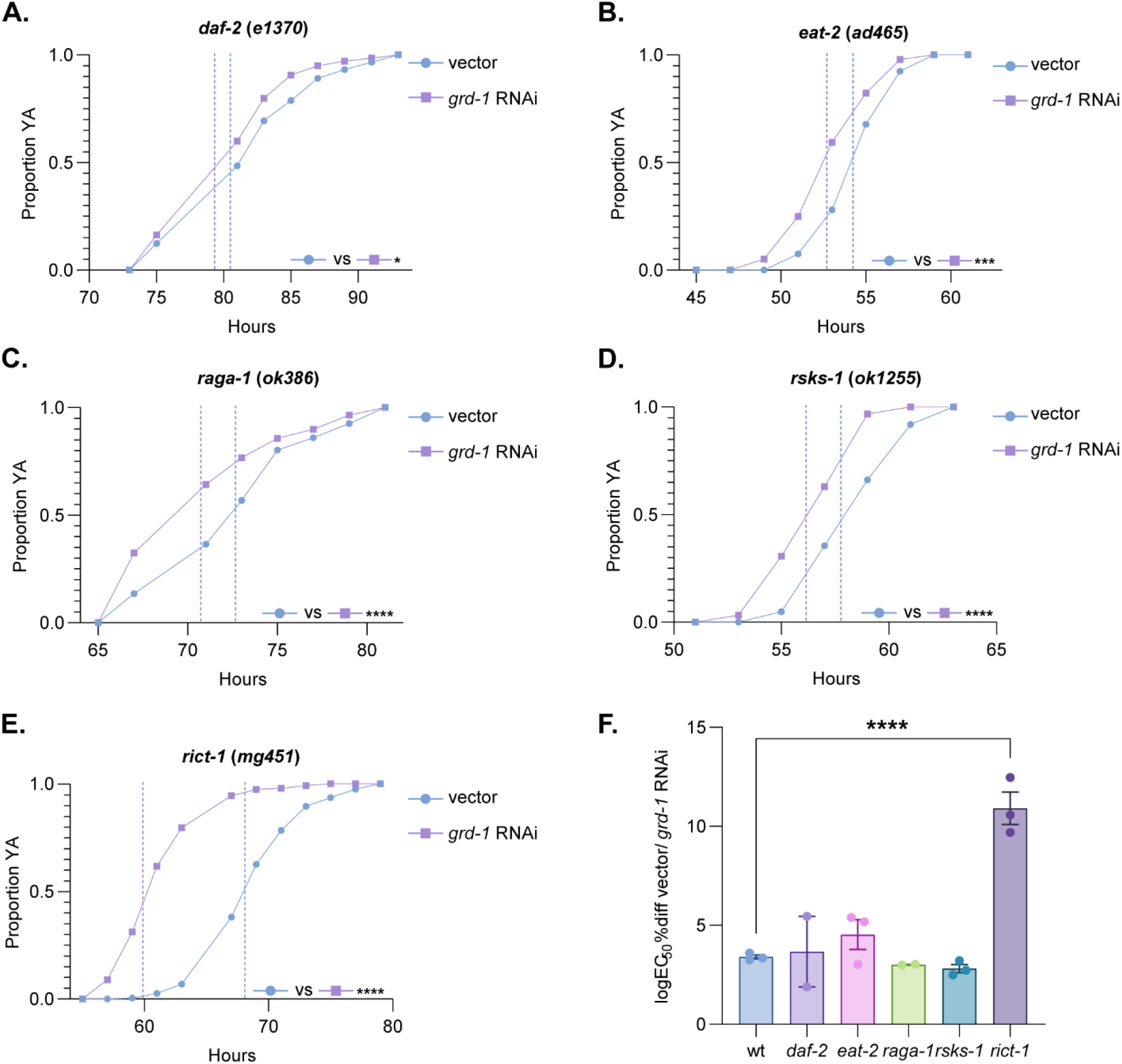
*grd-1* knockdown disproportionately accelerates development in TORC2 mutant *rict-1*, in contrast to other nutrient sensing pathways. (**A**-**D**) *grd-1* RNAi accelerates *daf-2, eat-2, raga-1*, and *rsks-1*, mutant developmental rates by ∼2 hours, quantitatively comparable to wild type (*n* > 50, \**P*<0.05, \*\*\**P*<0.001, 01, by log-rank test). (**E**) *grd-1* RNAi accelerates *rict-1* mutant development by ∼6 hours (Bonferroni *P*<0.0001 by log-rank test). (**F**) LogEC50 percent difference in hypothetical midpoint times in transition to adulthood between the vector and *grd-1* RNAi developmental curves for each mutant described in (**A**) to (**E**) shows that the difference between vector and *grd-1* RNAi developmental curves for *rict-1* mutants is significantly greater than for wild type and other mutants tested (*n* ≥ 2, *P*<0.0001 by one-way ANOVA with Dunnett’s correction for multiple comparisons). Dashed lines represent hypothetical midpoint times at which 50% of the population has reached young adulthood. Data represented as mean ± s.e.m. Data from a minimum of two biological replicates shown in supplementary Table S1.

### *grd-1* knockdown rescues slowed growth in TORC2 pathway *sgk-1* and *sinh-1* loss of function mutants, and *grd-1* overexpression is sufficient to slow growth

Previous work has identified the serine-threonine kinase SGK-1 as a major downstream effector of TORC2 in *C. elegans* and *Saccharomyces cerevisiae* (Aronova et al., 2008; Jones et al., 2009; Webster et al., 2013; Zhou et al., 2019). *sgk-1* loss-of-function mutants phenocopy *rict-1* mutants’ slowed development, reduced brood, small body size, and shortened lifespan (Webster et al., 2013; Zhou et al., 2019). We therefore tested whether *grd-1* suppression functions to accelerate *rict-1* hypomorph development by increasing SGK-1 activity or by acting downstream of *sgk-1*. We find that *sgk-1* mutant development on *grd-1* RNAi is accelerated by ∼6 hours, similar to *rict-1* mutants (Fig. 3A; Table S1). Further substantiating that *grd-1* acts downstream of both *rict-1* and *sgk-1, grd-1* knockdown on a *rict-1*;*sgk-1* double mutant speeds up development by ∼6 hours (Fig. 3B; Table S1). Slowed development caused by lowered TORC2 signaling in loss-of-function mutants in the essential TORC2 subunit *sinh-1*(Sin1) is similarly accelerated by *grd-1* knockdown (Fig. 3C; Table S1). Finally, broader relevance of *grd-1* to TORC2 biology is indicated by the fact that *grd-1* RNAi also rescues shortened lifespan, increased fat mass, small body size, and reduced brood size of *rict-1* mutants (Fig. S2A-D; Table S3) (Soukas et al., 2009).

**Fig. 3.**
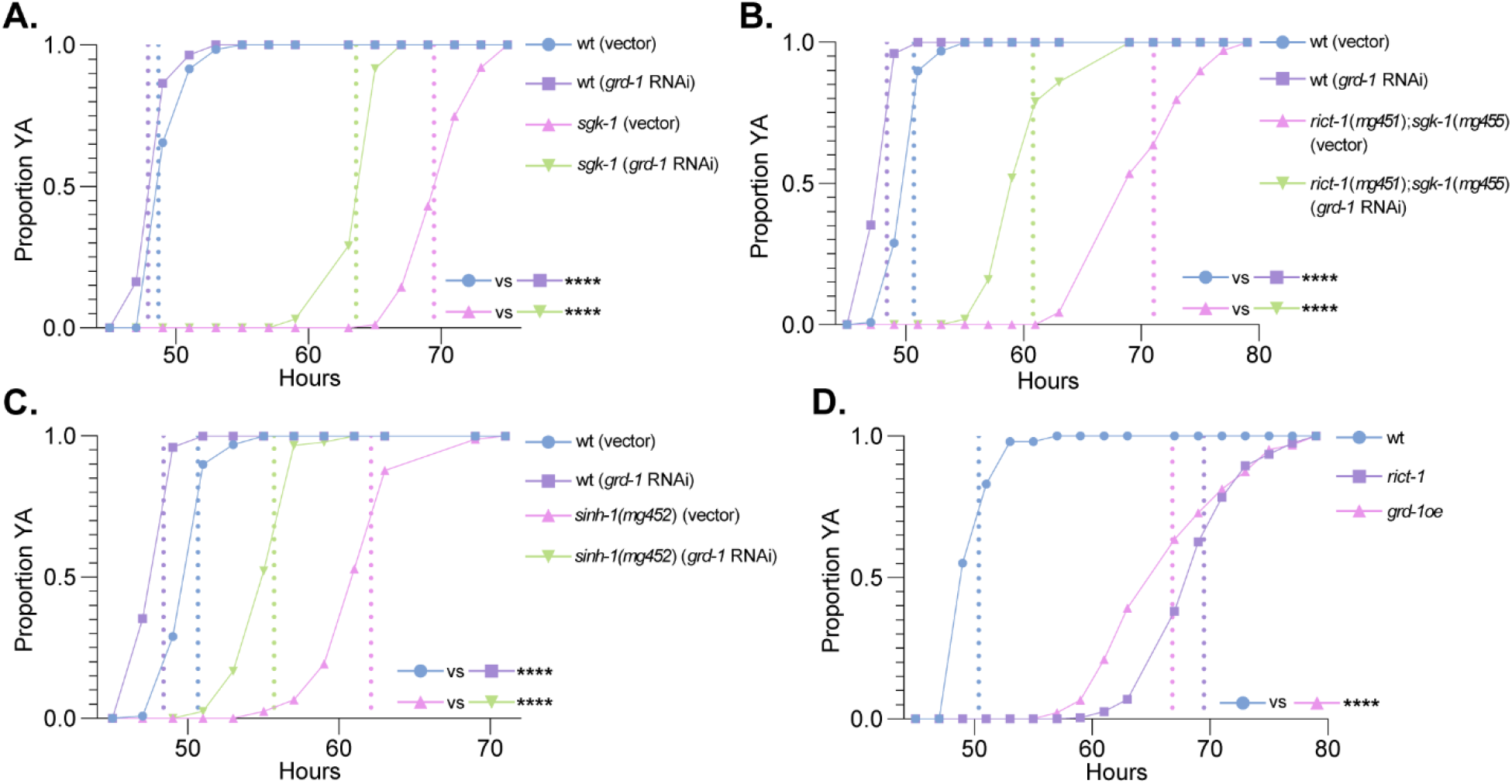
*grd-1* knockdown disproportionately accelerates development in multiple TORC2 signaling defective backgrounds and overexpression is sufficient to slow development. (**A**) Developmental rate of *sgk-1*(*mg455*) mutants is accelerated by ∼6 hours by *grd-1* RNAi (*n* > 100, *P*<0.0001 by Bonferroni corrected log-rank test). (**B**) Developmental rate of *rict-1*(*mg451*);*sgk-1*(*mg455*) double mutants is accelerated by ∼6 hours by *grd-1* RNAi (*n* > 50, *P*<0.0001 by Bonferroni corrected log-rank test). (**C**) Developmental rate of *sinh-1*(*mg452*) mutants is accelerated by ∼6 hours by *grd-1* RNAi (*n* > 50, *P*<0.0001 by Bonferroni corrected log-rank test). (**D**) *grd-1* overexpression is sufficient to slow developmental rate similar to *rict-1* mutant developmental rate (*n* > 50, *P*<0.001 by Bonferroni corrected log-rank test). Dashed lines represent hypothetical midpoint times at which 50% of the population has reached young adulthood. Data in biological triplicate shown in supplementary Table S1. Note the same N2 controls are used in B and C.

As *grd-1* is necessary for slowed development downstream of TORC2, this suggests that increased *grd-1* activity might be sufficient to slow development. In order to test this possibility, we generated transgenic *C. elegans* overexpressing *grd-1* under the control of its native promoter. Indeed, *grd-1* overexpression is sufficient to prompt slowing of developmental rate of wild type animals to a degree similar to *rict-1* mutants (Fig. 3D).

We have previously identified non-canonical activity of the dosage compensation complex (DCC) as partially necessary for developmental delay downstream of TORC2 (Webster et al., 2013). This raised the possibility that *grd-1* may be acting downstream of or in concert with the DCC. However, *grd-1* transcript levels are unperturbed in both *rict-1* mutants and DCC subunit *dpy-21* knockdown by RNAi treated animals, suggesting that *grd-1* is not transcriptionally regulated by either TORC2 or the DCC during larval development (Fig. S3A-B).

### *ptr-11* knockdown rescues development in TORC2 mutants and in *grd-1* overexpressors

Thus, since *grd-1* is not transcriptionally regulated, we hypothesized that its activity is likely to be increased in TORC2 mutants. In order to begin to identify how what we hypothesize to be increased activity of *grd-1* may be transduced, we performed a screen on Patched and Patched-related receptor (*ptc* and *ptr*, respectively) gene families to identify phenocopiers of *grd-1*. Of *ptc* and *ptr* family members tested by RNAi, only *ptr-11* knockdown accelerates development (Fig. 4A). Indeed, treatment of *rict-1* mutants, *rict-1*;*sgk-1* double mutants, and *sinh-1* mutants with *ptr-11* RNAi produces quantitatively similar acceleration of growth rate as *grd-1* RNAi (Fig. 4B; Fig S4A-B; Table S1), suggesting that *grd-1* and *ptr-11* may function in a common genetic pathway downstream of TORC2/SGK-1 signaling. Consistent with this possibility, in slow-growing *grd-1* overexpressors, *ptr-11* RNAi accelerates development by ∼4 hours compared to ∼1 hour in wild type animals, substantiating the notion that *ptr-11* functions downstream of *grd-1* (Fig. 4C; Fig. S4E; Table S1).

**Fig. 4.**
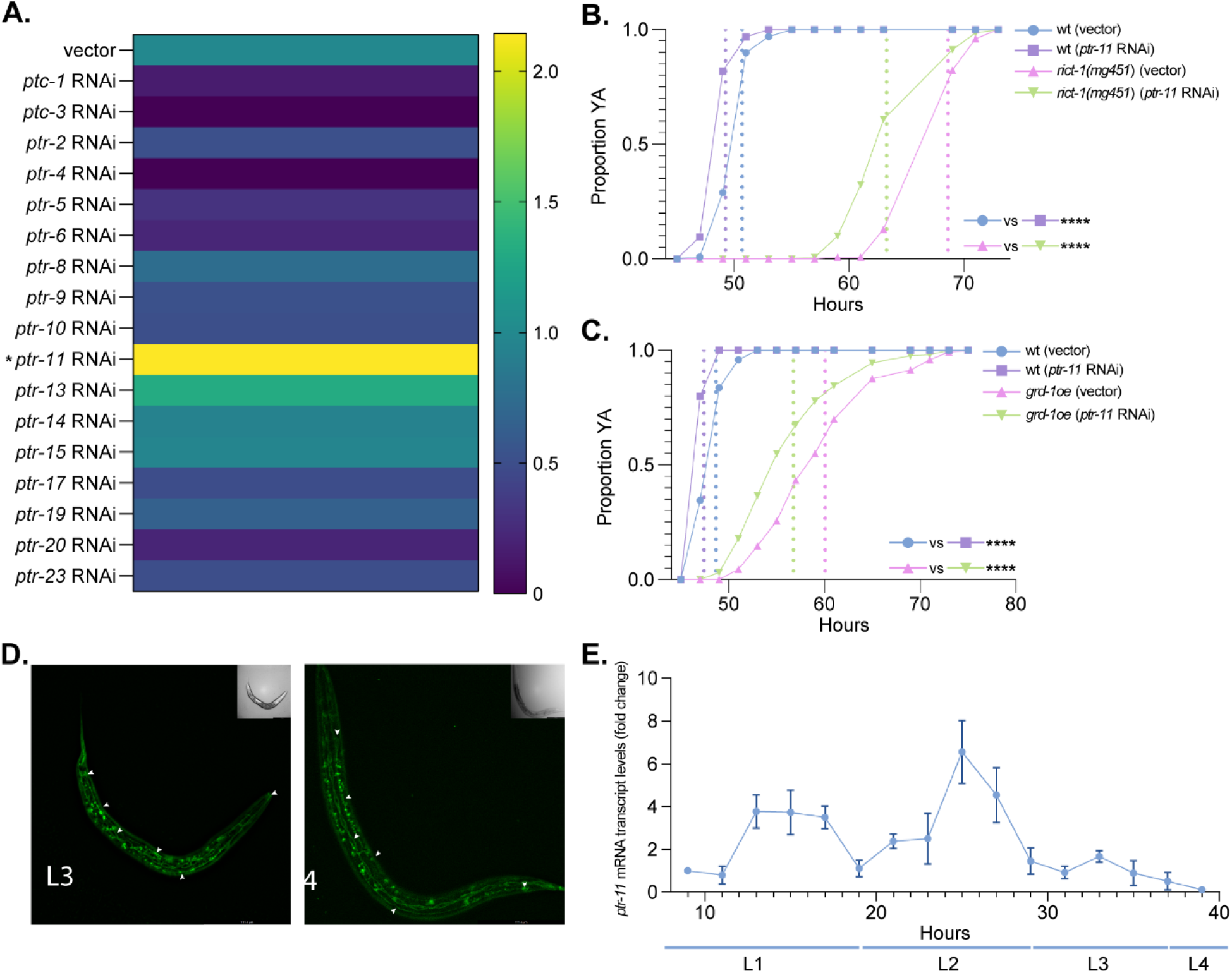
*ptr-11* knockdown phenocopies *grd-1* knockdown and significantly rescues slowed *grd-1oe* development. (**A**) Developmental screening of *ptc*/*ptr* family members reveals that *ptr-11* RNAi also accelerates wild type *C. elegans* development (*n* = 3, \**P*<0.1 by one-way ANOVA with two-stage Benjamini-Krieger-Yekutieli FDR-adjustment). Note that vector = 1. (**B**) *ptr-11* knockdown accelerates *rict-1*(*mg451*) development by ∼5 hours (*n* > 50, *P*<0.0001 by Bonferroni corrected log-rank test). (**C**) *ptr-11* RNAi significantly rescues slowed *grd-1oe* development (*n* > 50, *P*<0.0001 by Bonferroni corrected log-rank test). Dashed lines represent hypothetical midpoint times at which 50% of the population has reached young adulthood. Data in biological triplicate shown in supplementary Table S1. Note that the same wild type vector control used in (**B**) is used in Fig. S4A-B. (**D**) An endogenous *ptr-11*::*GFP* fusion shows expression in the hypodermis (where expression appears to be on the cell surface), dorsal and ventral nerve cords, neurons, and seam cells at both the L3 and L4 stages. (**E**) Similar to *grd-1, ptr-11* mRNA expression oscillates with molting.

Using an endogenously tagged *ptr-11*::*GFP* generated by CRISPR/Cas9 genome engineering, we observe *ptr-11* expression in the hypodermis, head and tail neurons, dorsal and ventral nerve cords, and seam cells at both the L3 and L4 stages (Fig. 4D). As is the case for *grd-1*, previous transcriptomics analyses have shown that *ptr-11* oscillates in expression throughout development (Hendriks et al., 2014), which we confirmed by qRT-PCR (Fig. 4E).

### *pqm-1* and its target genes are dysregulated in *rict-1* mutants in a *grd-1* dependent manner

In order to identify potential downstream effectors of *grd-1/ptr-11* signaling that mediate growth rate downstream of TORC2, we returned to our RNA sequencing results. Analysis of these data indicate that target genes of transcription factors PQM-1 and BLMP-1 are enriched following *grd-1* knockdown (Fig. 5A; Table S3). Assessment of developmental rate of both wild type and *rict-1* animals with *pqm-1* and *blmp-1* RNAi indicates that only *pqm-1* knockdown accelerates development (Fig 5B; Fig S5A). From our RNA sequencing data, we identified the six *pqm-1* target genes that are most decreased by *grd-1* suppression and find that two of these, *C49G7*.*7* and *ugt-43*, are upregulated in *rict-1* loss of function mutants and confirmed to be downregulated by *grd-1* RNAi by qRT-PCR (Fig. 5C). This suggests that PQM-1 activity is increased in TORC2 loss of function in a *grd-1-*dependent manner. In support of the conclusion that *grd-1* drives PQM-1 activity, *pqm-1* knockdown decreases *ugt-43* expression in *rict-1* mutants (Fig. S5B). Further, RNAi against *ugt-43* mildly accelerates development in both wild type and *rict-1* mutant animals (Fig 5D), indicating that targets of PQM-1 are indeed active in governance of developmental rate, albeit to a lesser degree than PQM-1 itself, as expected.

**Fig. 5.**
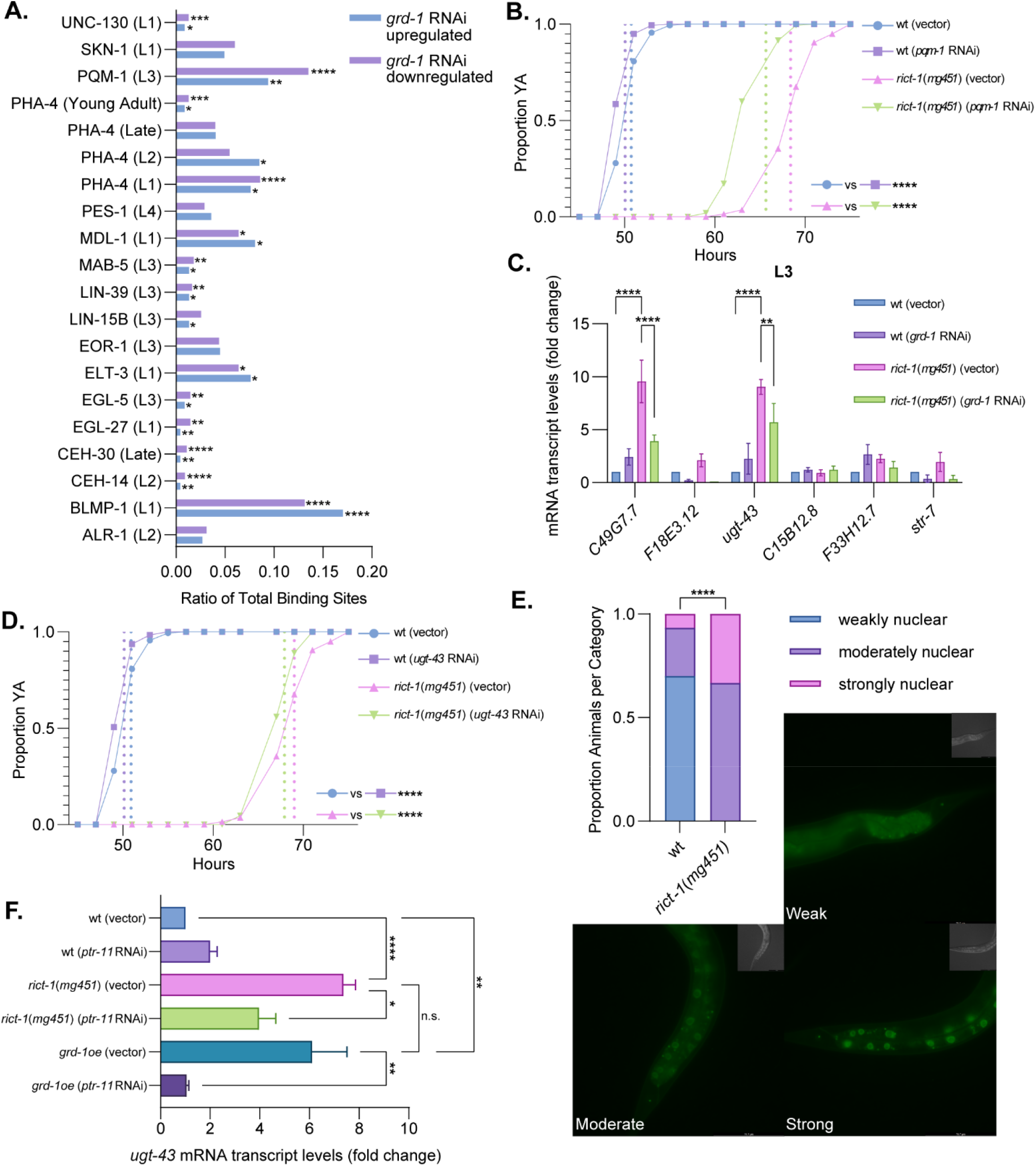
*pqm-1* knockdown partially phenocopies *grd-1* knockdown and *pqm-1* target genes dysregulated in *rict-1*(*mg451*) mutants are *grd-1* and *ptr-11* dependent. (**A**) Analysis of the ratio of total modENCODE transcription factor binding sites for differentially regulated transcripts in *grd-1* suppressed L3 animals reveals that PQM-1 and BLMP-1 are most positionally enriched (*n* = 3, adjusted two-tailed binomial \**P*<0.05, \*\**P*<0.01, \*\*\**P*<0.001, \*\*\*\**P*<0.0001). (**B**) *pqm-1* knockdown accelerates development of wild type animals by ∼1 hour and *rict-1*(*mg451*) mutant development by ∼4 hours (*n* > 100, \*\*\*\**P*<0.0001 by Bonferroni corrected log-rank test). (**C**) Measuring transcript levels for six genes most downregulated by *grd-1* RNAi reveals that both *C49G7*.*7* and *ugt-43* are upregulated in *rict-1* animals at the L3 stage and significantly downregulated by *grd-1* knockdown (*n* = 3, \*\**P*<0.01, \*\*\*\**P*<0.0001, by two-way ANOVA with Tukey’s correction for multiple comparisons). (**D**) *ugt-43* knockdown accelerates wild type development by ∼1 hour and *rict-1* development by ∼2 hours (*n* > 100, \*\*\*\**P*<0.0001 by Bonferroni corrected log-rank test). (**E**) *pqm-1*::*GFP* nuclear localization in the posterior intestine is significantly increased in *rict-1*(*mg451*) mutants relative to wild type animals (*n* = 10, three biological replicates, \*\*\*\**P*<0.0001 by Chi square goodness of fit test). Representative images for weak, moderate, and strong binning categories are shown on the right. (**F**) *ugt-43* mRNA transcript levels are upregulated in both *rict-1*(*mg451*) and *grd-1oe* animals and downregulated by *ptr-11* RNAi in both backgrounds (*n = 3*, \**P*<0.05,\*\**P*<0.01,\*\*\*\**P*<0.0001, by one-way ANOVA with Dunnett’s correction for multiple comparisons). Note *ugt-43* and *pqm-1* curves (B and D) are nested with the same control. Developmental data available in biological duplicate in supplementary Table S1. Data represented as mean ± s.e.m.

To further elucidate a connection between *rict-1* and *pqm-1*, we examined endogenously tagged *pqm-1*::*GFP* nuclear localization at the L3 stage in both wild type and *rict-1* loss of function mutants. PQM-1::GFP shows strong nuclear localization in *rict-1* mutants vs. wild type (Fig. 5E). Using *ugt-43* as a proxy of *pqm-1* activity, we further show that PQM-1 is more active at the L3 stage in both *rict-1* mutant and *grd-1* overexpressing animals (Fig 5F). In support of a TORC2/SGK/GRD-1/PTR-11/PQM-1 signaling axis, increased PQM-1 activity read out as *ugt-43* expression in TORC2/*rict-1* mutants and *grd-1* overexpression transgenics is dependent upon *ptr-11*, as knockdown abrogates increased *ugt-43* mRNA levels (Fig. 5F).

## DISCUSSION

Correctly integrating nutritional signals from the environment is paramount to ensuring optimal growth rate and development. Ours and others’ previous work indicates that TORC2 is critical governance of growth, reproduction, and lifespan, regulating these processes in a diet-dependent manner (Jones et al., 2009; Soukas et al., 2009). Defects in TORC2/SGK-1 signaling lead to slow growth, substantiating the important role of this complex in ensuring environmentally-appropriate organismal growth. In this report, we show that an effector arm of slowed growth downstream of TORC2 is a heretofore unappreciated *grd-1*/*ptr-11* Hedgehog signaling cascade that relies on chronic stress transcription factor *pqm-1*. Based upon data presented here, we suggest that increased *grd-1* activity signals through *ptr-11* to slow growth by activating *pqm-1* transcriptional responses. This work not only identifies a new cognate ligand-receptor pair of Hh-Ptr in *grd-1*/*ptr-11* but is also the first indication that TORC2 and Hh signaling cooperate to control whole organismal development, growth, and metabolism.

The study of developmental timing in *C. elegans* has most extensively explored heterochronic genes required for correct developmental event sequencing, such as *lin-4, lin-14*, and *let-7* (Ambros and Horvitz, 1984; Ambros and Moss, 1994; Reinhart et al., 2000). By comparison, there have been fewer dissections of the specific mechanisms controlling developmental rate in *C. elegans*. Many such investigations have focused on pathways controlling developmental progression in the absence of food or certain nutrients, revealing critical roles for IIS signaling, TORC1, lipid metabolism, the one-carbon cycle, and micronutrients in dauer formation and larval arrest (Baugh and Sternberg, 2006; Galles et al., 2018; Long et al., 2002; Watson et al., 2014; Watts et al., 2018). However, our understanding of the role of TORC2 signaling in development remains largely incomplete, confined to the DCC, a gut-neuronal axis linking TORC2 to TGF-β signaling, and CDC-42 induced neuronal protrusions (Alan et al., 2013; O’Donnell et al., 2018; Webster et al., 2013). Here, our results point to a mechanism by which Hh-r morphogens *grd-1*/*ptr-11* act downstream of TORC2 to control development and growth. Although study of *grd-1* and *ptr-11* activity *per se*, is challenging, and given that neither *grd-1* nor *ptr-11* are regulated at the RNA or protein level downstream of TORC2, we are able to connect TORC2 inactivation to increased GRD-1 signaling by invoking downstream activity of PQM-1. Specifically, both loss of TORC2 and *grd-1* overexpression lead to increased PQM-1 activity measured by target gene expression, suggesting that under conditions of lowered TORC2 signaling, activity of *grd-1*/*ptr-11* is induced to slow development. Although we find that these Hh-r proteins function downstream of major TORC2 effector kinase SGK-1, it remains unclear how *grd-1* activity is directly regulated. However, the lack of *grd-1* transcriptional changes in *rict-1* loss of function mutants suggests a post-transcriptional regulatory mechanism. Attempts to make an endogenously tagged mature GRD-1 protein were not successful and have proven challenging by previous reports (Aspöck et al., 1999), so additional work is needed to define the precise mechanisms of GRD-1 activation. Notably, our data do not show a transcriptional interaction between the DCC and *grd-1*, indicating that TORC2 likely controls development through these two arms in parallel.

*C. elegans* has a much-expanded family of Hedgehog morphogens that are thought to share a common ancestor with other phyla (Aspöck et al., 1999). Further, *C. elegans* possesses an expanded family of Patched-related receptors but lacks several canonical Hedgehog signaling components such as Smoothened and a truly orthologous Gli transcription factor (Zugasti et al., 2005). Hh-r morphogens and Ptr receptors have shared biological functions in controlling developmental progression and have been shown to interact cell non-autonomously with one another as in the case of *wrt-10*/*ptc-1* and *grl-21*/*ptr-24* (Lin and Wang, 2017; Templeman et al., 2020).

Here we provide just the third example of an Hh-r-Ptr pair that functions in tandem to regulate metabolism in *C. elegans*. Our findings complement prior reports indicating that Hh-r proteins control important parts of organismal homeostasis such as reproductive health downstream of CREB via *wrt-10*/*ptr-2* signaling (Templeman et al., 2020). Based upon patterns of *grd-1* promoter activity, we suggest that *grd-1* is produced in the intestine in a cyclical fashion during molting, and that under conditions of lowered TORC2 signaling, *grd-1* function is increased to put the brakes on development. When *grd-1* levels are artificially increased by overexpressing the protein under its native promoter, worm development is slowed to near *rict-1* hypomorph levels. Given that the intestine is both the primary tissue where *grd-1* is expressed and the principal tissue of action for TORC2 in regulation of growth and metabolism (Soukas et al., 2009), it seems plausible that TORC2 controls development by modulating *grd-1* activity in the intestine. However, like other Hh-r proteins, *grd-1* is predicted to be an extracellular protein, suggesting a cell non-autonomous role for the morphogen. We identify *ptr-11* as the likely receptor for *grd-1* given that its knockdown significantly restores developmental rate in TORC2 mutants and *grd-1* overexpression transgenics. Our endogenous *ptr-11*::*GFP* fusion indicates cell surface expression in several tissues including the hypodermis and seam cells, but not the intestine. Definitive determination of the respective sites of action of *grd-1* and *ptr-11* will require additional investigation. In aggregate, these results compellingly suggest that TORC2/SGK-1 takes stock of nutrient and growth environment status. If conditions are favorable, TORC2/SGK-1 signaling tamps down *grd-1* activity. However, in response to unfavorable conditions, reduced TORC2/SGK-1 signaling increases *grd-1* activity, likely in the intestine, and *grd-1* executes a program delaying organismal growth by relaying the signal of unfavorable conditions to other tissues through *ptr-11*.

Moreover, we identify *pqm-1* as a downstream effector of the *grd-1*/*ptr-11* signaling relay. Globally, PQM-1 acts antagonistically to DAF-16 in IIS signaling mutants to regulate lifespan and development (Tepper et al., 2013). Also, PQM-1 has been previously shown to work downstream of TORC2 to mobilize fat from the intestine to the germline at the onset of adulthood (Dowen et al., 2016). Although *pqm-1* loss of function mutations cause slight developmental delay (Tepper et al., 2013), larval knockdown by RNAi causes developmental acceleration that partially recapitulates *grd-1*/*ptr-11* knockdown in both wild type animals and TORC2 hypomorphs. Using increased expression of *pqm-1* target gene *ugt-43* as a barometer for increased *pqm-1* activity in *rict-1* and *grd-1oe* animals, we conclude that *grd-1* is activated in *rict-1* mutants, which in turn activates *pqm-1* to effectuate a slowing of growth in a *ptr-11* dependent manner. These data strongly support our conclusion that under lowered TORC2 signaling conditions, growth is slowed by increased signaling through a *grd-1/ptr-11/pqm-1* signaling relay. Definitive proof that *grd-1* and *ptr-11* represent a ligand-receptor pair and how *ptr-11* signaling increases *pqm-1* activity will require more investigation.

In summary, we define a heretofore unappreciated coordination of the TORC2 and Hedgehog pathways in a signaling axis that functions to put the brakes on development when reduced TORC2 signaling indicates unfavorable environmental conditions. Future work will focus on which aspects of the nutrient milieu are sensed by TORC2 and how changes that prompt developmental slowing are communicated through the GRD-1/PTR-11/PQM-1 signaling relay delineated herein. Better understanding of this biology will inform how organisms govern growth rate through ancient and complex communication between diverse signaling networks.

---

**Fig. S1.**
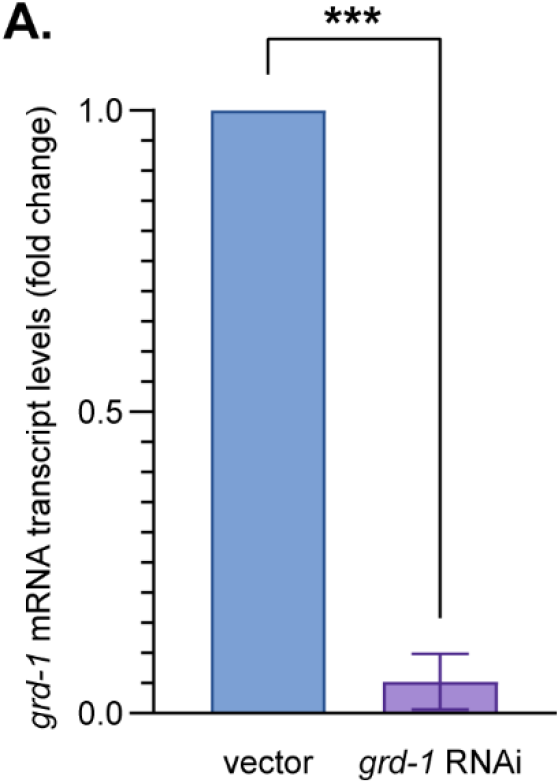
*grd-1* RNAi reduces *grd-1* mRNA transcript levels by 90%. (**A**) We confirm that the Ahringer library *grd-1* RNAi clone knocks down *grd-1* (*n = 3*, \*\*\**P*<0.001 by two-tailed Student’s T-test).

**Fig. S2.**
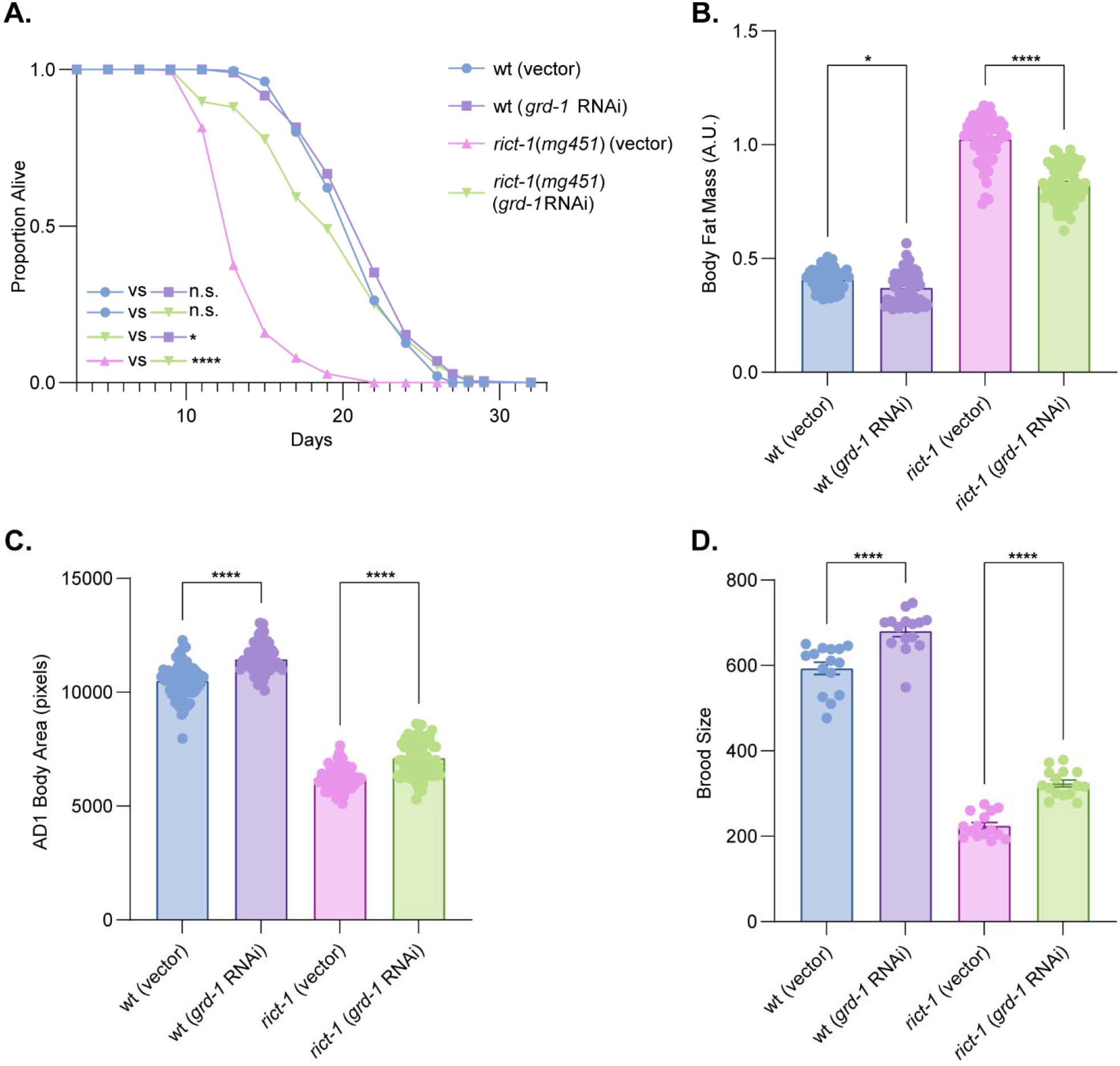
*grd-1* knockdown rescues multiple *rict-1*(*mg451*) healthspan pleiotropies. (**A**) *rict-1*(*mg451*) shortened lifespan is rescued to near wild type by *grd-1* RNAi (*n* > 100, *P*<0.0001 for wt vs. *rict-1*[vector] curves, *P*<0.0001 for *rict-1*[vector] vs. *rict-1*[*grd-1* RNAi], *P*<0.05 for wt[*grd-1* RNAi] vs. *rict-1*[*grd-1* RNAi], all other curves n.s. by Bonferroni corrected log-rank test). Data in biological triplicate shown in supplementary Table S3. (**B**) Body fat mass, as assessed by Nile red fixative staining, is reduced by *grd-1* RNAi in both wild type and *rict-1*(*mg451*) mutants (*n* > 15). (**C**) Body area is increased by *grd-1* RNAi in wild type worms and in *rict-1*(*mg451*) mutants. (**D**) Brood size is increased by *grd-1* RNAi in both wild type and *rict-1*(*mg451*) mutants (*n* = 5). \**P*<0.05, \*\*\*\**P*<0.0001 by two-way ANOVA with Tukey’s correction for multiple comparisons. Data from three biological replicates. Data represented as mean ± s.e.m.

**Fig. S3.**
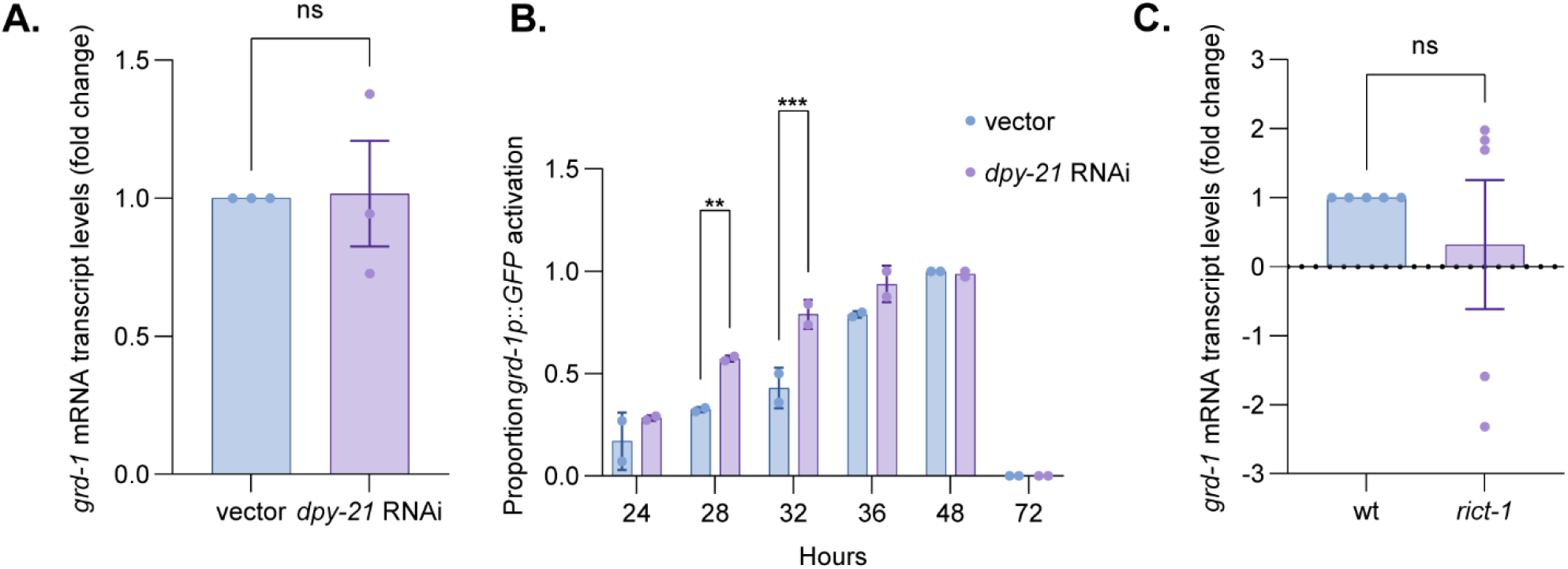
*grd-1* is not transcriptionally regulated in either *dpy-21* RNAi treated animals or *rict-1*(*mg451*) mutants. (**A**) *grd-1* mRNA levels are not significantly changed following *dpy-21* knockdown in wild type animals (*n* = 3, n.s. = non-significant, by two-tailed Student’s t-test). (**B**) Intestinal *grd-1* expression as measured by GFP expression is not downregulated in *dpy-21* RNAi (the hypothesized direction if *dpy-21* were driving increases in *grd-1* in *rict-1* mutants) treated animals at any timepoint. In fact, *dpy-21* knockdown induces earlier induction of intestinal GFP expression in *grd-1p*::*GFP* transgenic worms as shown by an increased proportion of GFP positive worms at 28 and 32 hours (*n* = 2, \*\**P*<0.01, \*\*\**P*<0.001, by two-way ANOVA with Tukey’s correction for multiple comparisons). (**C**) *grd-*1 transcript levels are not significantly dysregulated in *rict-1*(*mg451*) mutants (*n* = 5, n.s. = non-significant, by two-tailed Student’s t-test). Data represented as mean ± s.e.m.

**Fig. S4.**
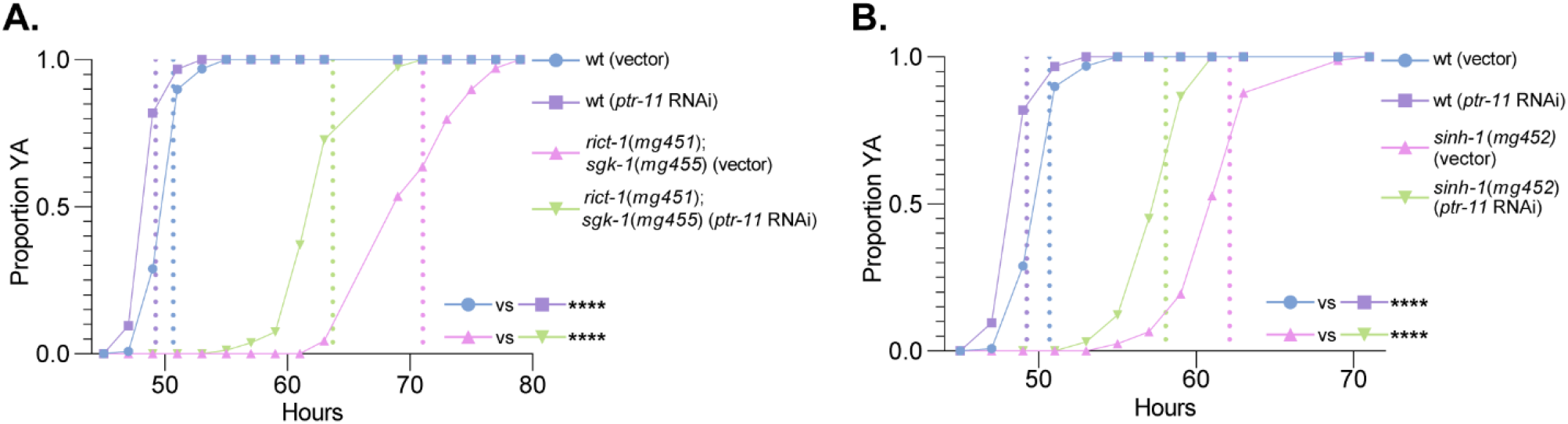
*ptr-11* RNAi significantly rescues development in other TORC2 hypomorphs. (**A**) *ptr-11* RNAi accelerates development of *rict-1*(*mg451*);*sgk-1*(*mg455*) mutants by ∼6 hours (*n* > 50, \*\*\*\**P*<0.0001 by Bonferroni corrected log-rank test). (**B**) *ptr-11* knockdown accelerates development of *sinh-1*(*mg452*) mutants by ∼4 hours (*n* > 50, \*\*\*\**P*<0.0001 by Bonferroni corrected log-rank test). Data available in biological triplicate in supplementary Table S1.

**Fig. S5.**
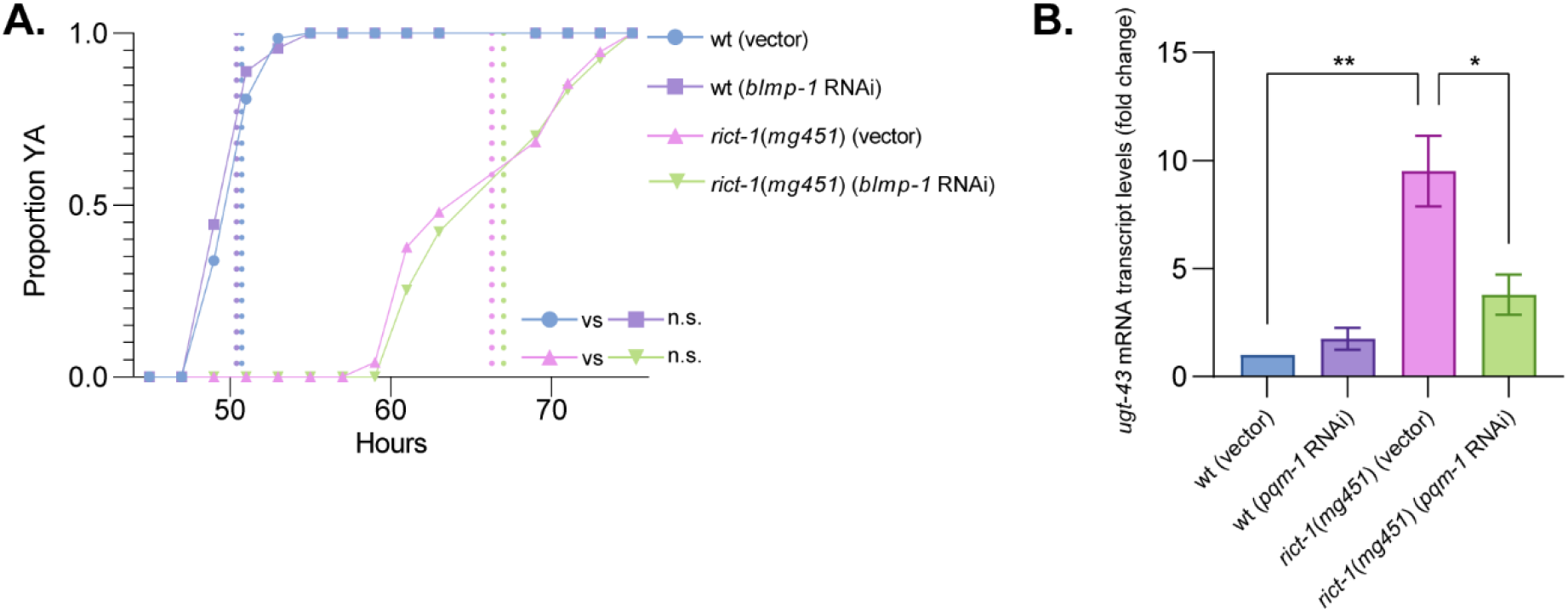
*blmp-1* RNAi does not accelerate development and *pqm-1* knockdown downregulates *ugt-43* expression. (**A**) *blmp-1* RNAi does not accelerate development in either wild type or *rict-1*(*mg451*) animals (*n* > 50, n.s. by Bonferroni corrected log-rank test). Data available in biological duplicate in supplementary Table S1. (**B**) *pqm-1* knockdown significantly reduces *ugt-43* mRNA transcript levels in *rict-1*(*mg451*) mutants (*n = 3*, \**P*<0.05, \*\**P*<0.01 by one-way ANOVA with Dunnett’s correction for multiple comparisons). Data represented as mean±s.e.m.

## Notes

### Competing Interest Statement

The authors have declared no competing interest.

